# Human macrophage polarization determines bacterial persistence of *Staphylococcus aureus* in a liver-on-chip-based infection model

**DOI:** 10.1101/2021.11.19.469246

**Authors:** Fatina Siwczak, Zoltan Cseresnyes, Swen Carlstedt, Anke Sigmund, Marko Gröger, Bas G.J. Surewaard, Oliver Werz, Marc Thilo Figge, Lorena Tuchscherr, Bettina Löffler, Alexander S. Mosig

## Abstract

Infections with *Staphylococcus aureus* (*S. aureus*) have been reported from various organs ranging from asymptomatic colonization to severe infections and sepsis associated with multiple organ dysfunction. Although considered an extracellular pathogen, *S. aureus* can invade and persist in professional phagocytes such as monocytes and macrophages. Its capability to persist and manipulate phagocytes is considered a critical step to evade host antimicrobial reactions. For the first time we leveraged a human liver-on-chip model and tailored image analysis algorithms to demonstrate that *S. aureus* (USA300) specifically targets macrophages in the liver models as essential niche facilitating bacterial persistence and phenotype switching to small colony variants (SCVs). *In vitro* M2 polarization was found to favor SCV-formation and was associated with increased intracellular bacterial loads in macrophages, increased cell death, and impaired recruitment of circulating monocytes to sites of infection. These findings expand the knowledge about the role of liver macrophages in the course of systemic infection. Further, the results might be relevant for understanding infection mechanisms in patients with chronic liver disease such as fibrosis that display increased frequencies of M2 polarized liver macrophages and have a higher risk for developing chronic infections and relapsing bacteremia.

## Introduction

*Staphylococcus aureus* (*S. aureus*) is a gram-positive bacterium representing an important human pathogen in community and hospital-acquired infections. Invasive infections can be associated with multiple organ dysfunction and prolonged treatment requirements [1]. Further, antibiotic resistance of invasive methicillin-resistant *S. aureus* (MRSA) complicates treatment of *S. aureus* infections associated with a high mortality rate of approximately 20% [2]. Infections with *S. aureus* are thus a leading cause of death in developed countries [3], with the MRSA strain USA300 reaching pandemic status across North America [4]. To date, no approved vaccine for *S. aureus* infections is available [5].

Various cell types, including non-professional phagocytes (such as endothelial, epithelial cells and keratinocytes), and professional phagocytes (e.g., macrophages), have been reported to provide shelter for intracellular persisting *S. aureus* [6]. In particular, macrophages have been identified as a privileged environmental niche for *S. aureus* persistence, offering protection from antimicrobial activity and detection by immune cells. *S. aureus* has been reported to manipulate macrophage activation for its own survival [7].

With a blood perfusion rate of more than one liter per minute, the liver is not only among the most intensively perfused organs of the human body but also a central organ for filtering blood-borne infections as it harbors 80% of all tissue-resident macrophages (Kupffer cells, KCs) [8]. KCs are located within the sinusoids with optimal access to pathogens arriving in the liver. They are critical effectors of the early innate immune response by facilitating phagocytosis and killing bacteria from the bloodstream [8a]. KCs thus likely represent a crucial reservoir for intracellular persistence of *S. aureus* [9]. Indeed, 90% of *S. aureus* are sequestered from the blood by the liver and effectively killed by KCs [10], but a small fraction of these bacteria manage to survive intracellularly. Macrophages have been shown to have a finite capacity for intracellular killing of *S. aureus* when exposed to large inocula. The bacteria can then persist intracellularly in vacuoles of macrophages for several days before escaping into the cytoplasm and eventually causing host cell lysis [6d, 11]. Very few bacteria can escape from macrophages, which can be sufficient to establish a tissue infection abscesses [11b, 12]. In this context, it has been postulated that macrophages might represent an important bottleneck triggering a bacterial variant enrichment to improve the pathogens’ persistence and spread within the body subsequently. The formation of SCVs is part of the normal bacterial life cycle of *S. aureus*, i.e., as a response to starvation. However, the formation of SCVs has also been linked to antibiotic treatment, response to antimicrobial host peptides, and the pathogenesis of chronic and recurrent infections [13]. SCVs can revert from a quiescent metabolic state to the normal parental phenotype, which can induce recurrent infections [13-14]. The unstable SCV phenotype switching has been further linked to the development of antibiotic tolerance associated with a reduced bacterial growth rate [15]. The intracellular milieu has been thought of as an essential trigger of SCV formation [6a].

In a simplified model, macrophage activation has been categorized *in vitro* into the pro-inflammatory M1 and the anti-inflammatory M2 phenotype. M1 polarization is associated with the release of pro-inflammatory cytokines and mainly microbicidal activity as a primary antibacterial defense mechanism [16]. The M2 polarization has mainly been described under homeostatic conditions and in later stages of infection. It is associated with an anti-inflammatory profile and poor microbicidal capacity to restrain inflammation and induce tissue repair [17]. To elucidate the role of macrophage polarization in the liver on persistence, SCV formation, and dissemination of *S. aureus*, we leveraged a recently developed *in vitro* model of the human liver. The liver-on-chip is composed of endothelial cells, hepatocytes, and macrophages, enabling studies on the mutual interaction between the pathogen and most essential cell types of the liver (Figure 1). Our studies revealed that *S. aureus* (strain USA300) could specifically target macrophages and subvert them as an essential niche for bacterial persistence, replication, and phenotype switching. We demonstrate that M2 macrophage polarization favors SCV phenotype formation that is associated with increased intracellular bacterial loads. Interestingly, the SCV formation was not observed in mono-cell cultures of macrophages nor liver-on-chip without cocultured macrophages. 48h post-infection (p.i.), we found an increased number of bacteria persisting in interleukin-4 polarized M2 macrophages compared to non-activated (M0) or interferon-activated M1 macrophages. The increased bacterial burden in M2 macrophage was associated with increased cell death, a drop in albumin synthesis by hepatocytes, and impaired replenishment of macrophages through disturbed recruitment of circulating monocytes.

**Figure 1.**
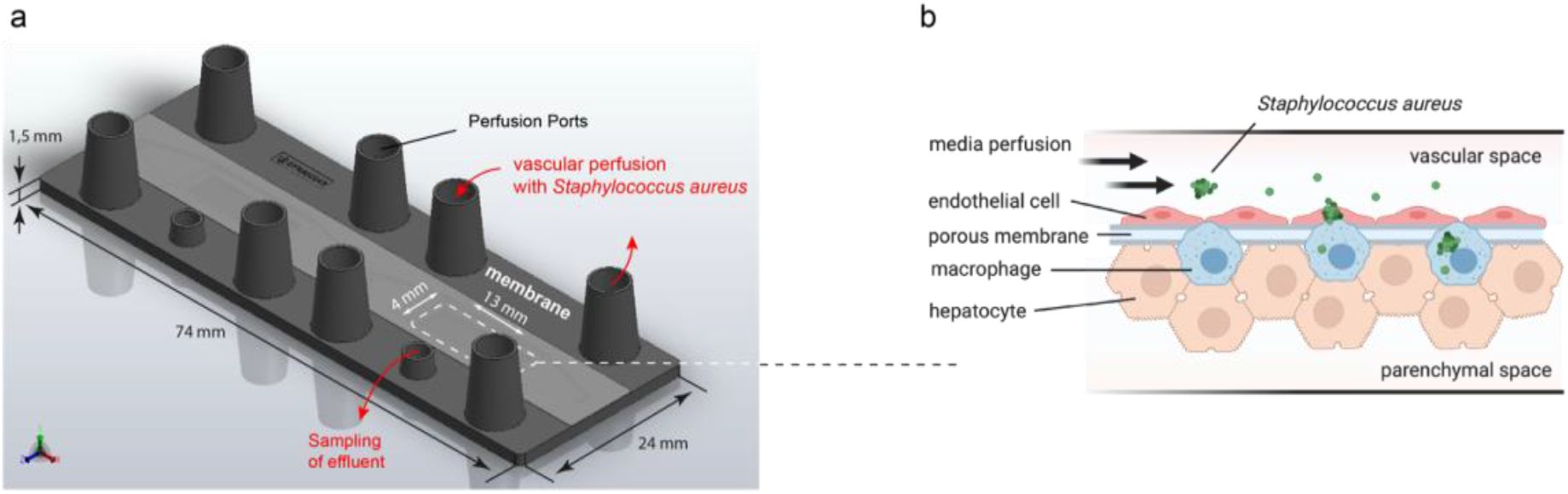
Drawing of the microfluidic perfused biochip for the culture of liver-on-chip. a) The microfluidic perfused chip has the format of a microscopic slide. It contains perfusion ports for the circulating medium to provide nutrition to cells cultured at the membrane suspended in each of the two cavities of the chip as recently described [18]. The effluent can be sampled under perfusion conditions via a sampling port. b) Scheme of the assembled cell layers in the liver on-chip. Endothelial cells are cultured at the top of the membrane and perfused by cell culture medium. At the opposite side of the membrane, hepatocytes are cultured. Macrophages are localized at the interface between the endothelial and the parenchymal cell layer formed by hepatocytes, as recently described [21]. For infection experiments, planktonic S. aureus were suspended in the cell culture medium perfused within the vascular space at the endothelial side of the liver-on-chip model.

## Materials and Methods

### Biochips

Microfluidic biochips were made from polybutylenterephthalat (PBT) and obtained from Dynamic42 GmbH (Jena, Germany). Biochips were manufactured by injection molding as described elsewhere [18]. The chamber above the membrane has a height of 700 μm; the chamber under the membrane has a height of 400 μm. The width of the afferent and efferent channels is 0.8 mm and 2 mm, respectively. The height of these channels is 0.6 mm and 0.4 mm, respectively. The upper and the lower chamber, including channel systems, have a volume of 220 μl and 120 μl, respectively. A 12 μm thin polyethylene terephthalate (PET) membrane with a pore diameter of 8 μm and a pore density of 1×10^5^ pores/cm^2^ (TRAKETCH Sabeu, Radeberg, Germany) was integrated. An area of 1.1 cm^2^ is available for cell culture. Chips and channels structures were sealed on top and bottom side with an extruded 140 μm thin Polystyrene (PS) foil using a laser welding-based proprietary bonding method. Gas permeable silicon tubing was used for perfusion, allowing oxygen equilibration during experiments. Ramping structures have been introduced into the chip bulk to prevent unfavorable flow conditions and trapping of stationary air bubbles. The biochip was perfused using peristaltic pumps (Ismatec REGLO digital MS-CA-4/12-100, Wertheim, Germany)

### Ethics

The study was approved by the ethics committee of the Jena University Hospital (2020– 1684, 3939-12/13), and all donors were informed about the aim of the study and gave written consent.

### Cell culture

Human umbilical vein endothelial cells (HUVECs) were isolated from human umbilical cord veins as described previously [19], cultured at a density of 3×10^5^ HUVEC’s/cm^2^ in endothelial cell growth medium (Endothelial Cell Medium, PromoCell, Heidelberg, Germany) with supplement. Donors were informed about the aim of the study and gave written consent. HUVECs were used until passage two. HepaRG cells were obtained from Biopredic International (Rennes, France). Cells were seeded at a density of 2.7×10^4^ cells/cm^2^ and cultured in William’s Medium E (Biochrom, Berlin, Germany) containing 10% (v/v) FCS (Life Technologies, Darmstadt, Germany), 5 mg/ml insulin (Sigma Aldrich, Steinheim, Germany), 2 mM glutamine (GIBCO, Darmstadt, Germany), 50 μM hydrocortisone-hemisuccinate (Sigma-Aldrich) and 100 U/ml Penicillin/100 mg/ml Streptomycin mixture (Pen/Strep) (GIBCO). The cells were cultured in a humidified cell incubator at 5% CO_2_ and 37°C for 14 days before differentiation. The medium was renewed every 3-4 days. Cell differentiation was induced as described [20], and cells were used for up to 4 weeks. Monocyte-derived macrophages were isolated as described previously [21]. Briefly, peripheral blood mononuclear cells (PBMCs) were isolated by Ficoll density gradient centrifugation as described previously [22] and seeded at a density of 1×10^6^ cells/cm^2^ in X-VIVO 15 medium (Lonza, Cologne, Germany) supplemented with 10% (v/v) autologous human serum. After 3 h incubation in a humidified cell incubator at 5% CO_2_ and 37°C, the cells were washed twice with X-VIVO 15 medium. Adherent monocytes were cultivated for 24 h in X-VIVO 15 medium and subsequently cultured in the liver model.

### Liver-on-chip

Liver models were assembled by staggered seeding of vascular and hepatic cell layers. In each sterilized biochip, 2.7×10^5^ HUVECs/cm^2^ and 1×10^5^ monocytes/cm^2^ were mixed and seeded on top of the membrane. HUVEC/monocytes were cocultured for five days with a daily medium exchange with vascular perfusion medium: endothelial cell medium supplemented with 20% autologous human serum, 10 ng/ml human granulocyte-macrophage colony-stimulation factor (GM-CSF) (Peprotech), and 10 ng/ml human macrophage colony-stimulating factor (M-CSF) (Peprotech) to induce macrophage differentiation. Subsequently, 2.7×10^5^/cm^2^ HepaRG were seeded on the membrane opposite to endothelial cells and macrophages and the HepaRG containing chamber filled with William’s E Medium (Biochrom) supplemented with 2 mM glutamine (GIBCO), Insulin (Sigma-Aldrich), hydrocortisone-hemisuccinate 50 μM (Sigma-Aldrich), 10% autologous serum. The chip was placed with HepaRG facing upward overnight to facilitate cell adhesion to the membrane. After one day of HepaRG attachment, the chip was flipped back, and perfusion was started at the vascular cavity with a flow rate of 50 μl/min. Medium at the lower chamber containing HepaRG was changed to hepatic cultivation medium: William’s E Medium (Biochrom) containing 2 mM glutamine (GIBCO), 0.5 μg/ml insulin ((Sigma Aldrich), hydrocortisone-hemisuccinate 50 μM (Sigma-Aldrich), 10% autologous human serum and 0.1% DMSO. The hepatic chamber was not perfused but medium was renewed daily.

### Macrophage polarization

Macrophages were allowed to differentiate within the endothelial layer for five days post-seeding. Macrophage polarization to M1 was induced by stimulation with 10 ng/ml IFNγ (Peprotech) or to M2 by stimulation with 10 ng/ml IL-4 (Peprotech) for two days. Polarization was performed by supplementing cytokines to the vascular perfusion medium, which was replaced by cytokine-free media immediately before infection (Supplementary Figure 1a).

### S. aureus culture

On the day of infection with living *S. aureus*, 1 ml of bacteria suspension from overnight culture was transferred to a 9 ml fresh MH medium containing 10μg/ml chloramphenicol and incubated at 37°C shaking for 3 h. Bacteria were pelleted and resuspended in PBS and adjusted to an optical density of 0.01. The number of bacteria determined by colony-forming unit (CFU) assays before and in parallel to each experiment in biological duplicates & technical triplicates. The multiplicity of infection (MOI) was calculated as the number of bacteria per cultured eukaryotic cell.

### Infection of liver-on-chip

Infection was performed at the vascular side of the liver-on-chip model with an MOI = 5 (referenced to endothelial cells and macrophages). Briefly, liver-on-chip devices were perfused with vascular perfusion medium containing bacteria for 90 min at 50 μl/min flow rate. Subsequently, extracellular bacteria were killed and removed by treatment with 20 μg/ml lysostaphin (WAK-Chemie Medical GmbH, Steinbach/Ts., Germany) for 30 min at the vascular and hepatic side of the model. Lysostaphin was washed away with vascular perfusion medium and hepatic cultivation medium, respectively. Infection was maintained for the indicated time points and flow conditions (50 μl/ml vascular perfusion). Vascular perfusion was stopped less than 5 min once a day for hepatic culture media exchange. Supernatants for CFU analysis were sampled at all indicated time points from the vascular perfusion stream and the hepatic chamber cultured under static conditions.

To analyze intracellular persisting bacteria, lysostaphin treatment was performed immediately before all cells of the liver-on-chip were lysed. Briefly, the membrane of the liver-on-chip was cut out, rinsed twice with PBS, and incubated in 0.5 ml PBS containing 0.25% Trypsin/EDTA (GIBCO, Germany) for 30 min at 37°C and 5% CO_2_. Subsequently, 1 ml ddH_2_O was added, harshly mixed, and centrifuged for 10 min at 13,000xg. Pellets containing living bacteria were resuspended in 0.5 ml PBS and plated as triplicates in serial dilution. CFU were counted 24, 48, and 72 post-plating, and CFUs determined and analyzed for the formation of small colony variants (SCV).

### Cytokine profiling

According to the manufacturer’s instructions, cytokines were measured from effluent at the vascular side using multiplex bead-based immunoassays (LEGENDplex, BioLegend, SanDiego, CA, USA). All samples were measured in duplicates for every donor and experiment using a BD Canto II flow cytometer (BD, Heidelberg, Germany). Data analysis was performed with LEGENDplex™ data analysis software (BioLegend, SanDiego, CA, USA).

### Monocyte recruitment assay

Monocytes were perfused in the absence or presence of suspended bacteria at a flow rate of 50 μl/min for 1 h through the liver-on-chip. 2.5 × 10^5^ monocytes were labeled with CellTracker™ Orange CMTMR Dye (ThermoFisher Scientific) according to manufactures recommendations, washed with PBS, and resuspended in 3 ml vascular perfusion medium. Flow through containing non adhesive monocytes was collected and monocytes treated with Lysostaphine for 30min, washed, and plated for CFU assay.

### Immunofluorescence staining

The following antibodies were used for staining: rabbit CD68 (BD Bioscience), goat epithelial cadherin (E-cadherin) (Thermo Fisher Scientific); donkey immunoglobulin (Ig)G anti-mouse IgG-Cy3, donkey IgG F(ab`), anti-rabbit IgG-AF488 (all obtained from Dianova, Hamburg, Germany). The membrane containing cell layers were removed from the biochip by carefully moving a scalpel along the edge of the cavity. The membrane was rinsed twice with PBS containing Ca^2+^ and Mg^2+^ (Lonza) and subsequently fixed with pre-cooled methanol at -20 °C for 10 min. Cell layers were fixed with 4% PFA. Blocking and permeabilization were done with PBS with 3% normal donkey serum (Dianova) 0.1% saponin (Sigma-Aldrich, Germany) for 45 min at RT. Primary antibodies were incubated overnight. The second antibody and DAPI were incubated after carefully washing twice with PBS and once with PBS/0.1 % saponin and incubated for 30 min at RT. Cells were washed four times with PBS containing Ca^2+^ and Mg^2+^ and mounted in a fluorescent mounting medium (DAKO, CA, USA). Finally, images were recorded at an AXIO Observer Z1 fluorescence microscope with an Apotome 2 extension (Carl Zeiss AG, Jena, Germany).

### Image analysis

Initially, an optical sectioning processing was carried out using ZEN 2 blue software (Carl Zeiss AG, Oberkochen, Germany) to provide images for further analysis as Z-stacks in the native Zeiss image format “CZI “. The Huygens Professional software (SVI, Hilversum, Holland) was used to deconvolve the CZI Z-stack images, utilizing the spinning disk deconvolution module at 4 μm pinhole spacing (see the SVI guidelines for handling Apotome image data). The deconvolved image stacks were segmented and quantified in Imaris 9.5.0, 9.5.1. or 9.6.0 (Bitplane, Zürich, Switzerland) according to the workflow in Supplementary Figure 2. The macrophages were reconstructed from the membrane-specific volume label together with the corresponding E-cadherin signal (Figure 2a-b). To segment the endothelial cells and the hepatocytes, the membrane locations were revealed by intensity-thresholding the E-, VE-cadherin and ZO-1 volume signals (Figure 2a-b). The membranes were reconstructed both as Surfaces (Figure 3b) and Cells objects (Figure 3c; for details on how the parameters were set up in Imaris for the membrane (Surfaces objects)- and cell-based (Cells objects) segmentation, see [23]). For the Cells objects, the DAPI channel served as guidance for the Nucleus component (Supplementary Figure 2); at the same time, the Cells objects were also reconstructed without nuclei in cases where the DAPI labeling was either not available or not necessary (Figure 2c). From the Cells objects, the membrane component was exported out individually, in order to be able to use them as Surfaces for masking and for counting the bacterial content in and/or near the endo- or epithelial cells (Figure 2c). The bacteria were exported out of the Cells module as “Vesicles” that were saved as Spots for further analysis (e.g., neighborhood density measurements).

**Figure 2.**
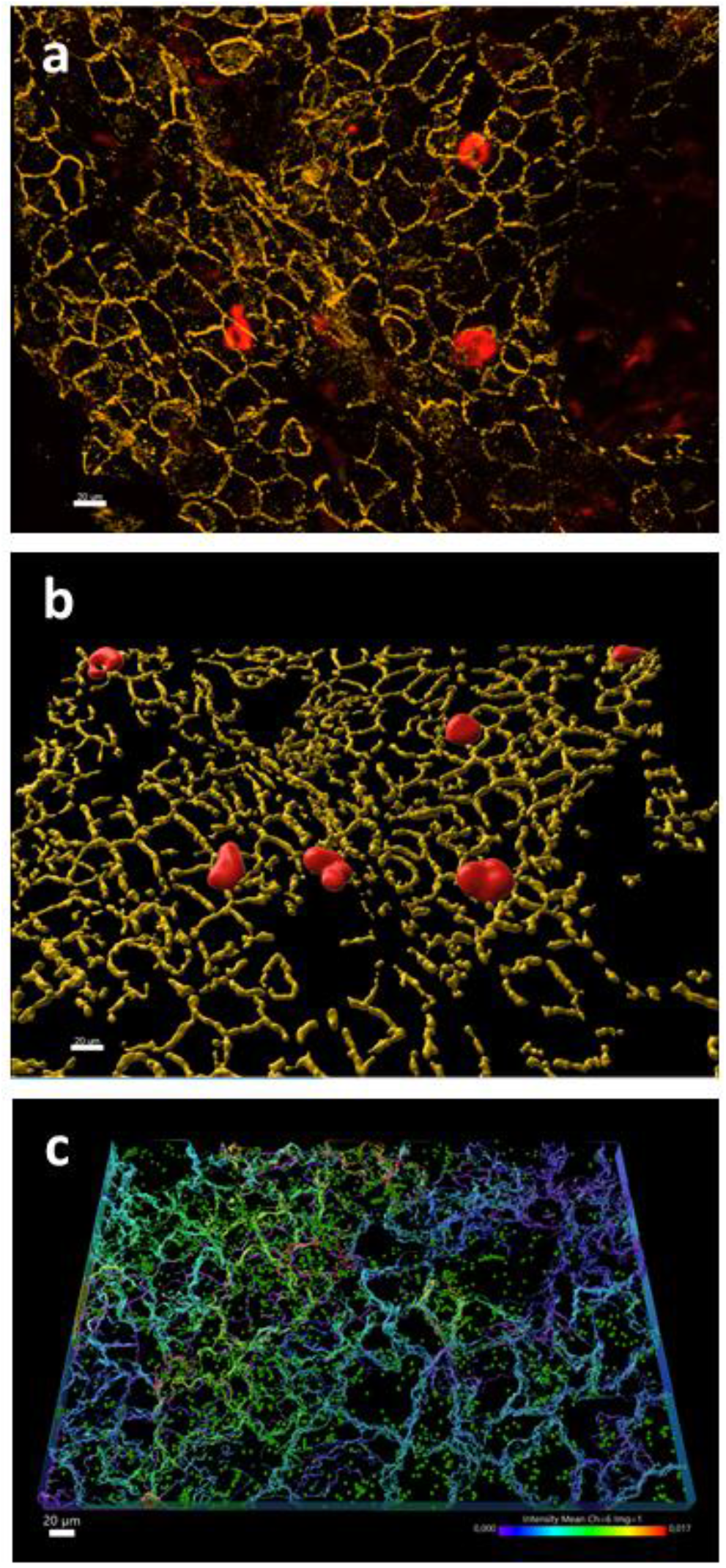
Illustrative processing steps of quantifying hepatocytes, macrophages, and S. aureus. The preprocessed (see Supplementary Figure 2) volume signals of the VE-cadherin (yellow) and macrophage (red) fluorescence channels **(a)** were segmented in Imaris, resulting in Surface objects in the corresponding colors (yellow: hepatocytes, red: liver macrophages) **(b)**. The VE-cadherin labelled cells and the S aureus bacteria were also segmented as Cells objects (see Supplementary Figure 2), followed by exporting the membrane components (as Cells) and the bacteria (as Vesicles) from the Cells objects **(c)**. The color coding of the cell membranes in (c) indicates the intensity of the green S. aureus fluorescence channel inside each individual endothelial cell (color scalebar in bottom right), thus characterizing the bacterial content of each cell. The S. aureus cells appear as green spots. The scalebars indicate 20 μm throughout.

**Figure 3.**
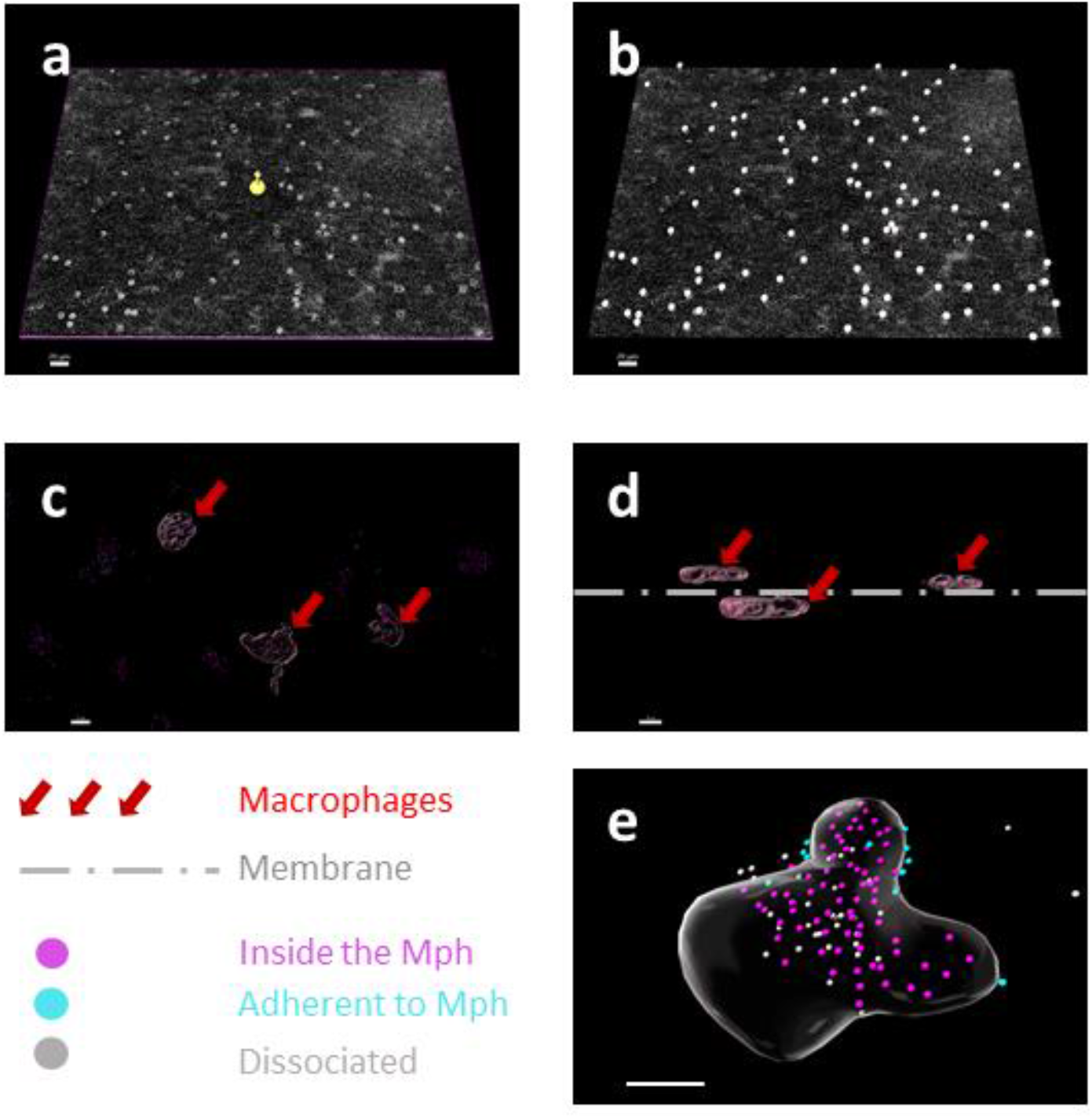
Identifying the relative spatial localization of immune cells and bacteria. **(a)** The green fluorescence channel was inverted (see also Supplementary Figure 2), contrast stretched and viewed in Slice mode in Imaris. The Z layer shown in (a) corresponds to the approximate middle of the membrane pores (bright disks). The yellow object in the center of (a) indicates the Imaris Z-slicing tool. **(b)** The bright disks were segmented as Spots after filtering by intensity and size (a pore is 8 μm in diameter) and were used to describe the location of each membrane pore (silver spheres). **(c)** Three macrophages were identified and segmented as Surfaces objects. **(d)** Their axial location was determined by comparing their Z position with that of the identified membrane pores (grey dashed-dotted horizontal line). The sideview shows that two of the macrophages were on the parenchymal side, whereas one was on the endothelial side of the membrane. **(e)** The locations of S. aureus cells relative to a macrophage are indicated by the color of the small spheres that illustrate the bacteria. The magenta spheres are bacteria that are inside the macrophage (the macrophage membrane appears as a grey glassy surface), the cyan spheres are adherent bacteria, whereas the silver spheres are dissociated pathogens that are more than one micrometer away from the macrophage surface. The scalebars in (a) and (b) indicate 20 μm, those in (c) to (e) are 10 μm.

### Statistical analysis

Statistical analysis was performed using GraphPad Prism software version 9.12 (GraphPad Software, La Jolla, CA, USA). One-way ANOVA analysis was performed with Dunnett’
ss correction for multiple comparisons. Two-way ANOVA was performed with Tukey’s correction for multiple comparisons. The use of the respective statistical tests is indicated in the figure legends with p-values labeled at the individual statistically significant data points.

## Results

Recent studies demonstrated an important role of liver macrophages in resolving bloodstream infections with *S. aureus* and revealed the potential of *S. aureus* to subvert this cell type as a privileged environmental niche [11a]. Yet, the role of macrophage plasticity as a potential determinant in the course of *S. aureus* infection needs to be elucidated.

The role of macrophages in pathogen persistence and dissemination was characterized in liver-on-chip cocultured with non-activated human macrophages (M0) or macrophages stimulated by IFNγ creating an M1-like phenotype or with IL-4 inducing an M2-like phenotype (Figure 1, Supp. Fig. 1a). After 2 h post-infection (p.i.) we analyzed intracellular bacterial counts in endothelial cells, hepatocytes, and macrophages using fluorescence microscopy and automated image analysis (Supp. Fig. 2). Briefly, a custom-designed algorithm was implemented in the Imaris software to preprocess and segment Z-stacks of microscopy images based on fluorescence intensity. The cells (macrophages, endothelial cells, hepatocytes) were identified as 3D surfaces using fluorescence membrane labelling, whereas the bacteria were characterized as uniformly sized spheres defined by their intensity center. The relative location of the cells (endothelial vs. parenchymal) and of the bacteria (inside vs. outside vs. adherent to macrophages or endothelial cells and hepatocytes) were quantified as described in the Methods section (Figures 2 and 3). The E-cadherin and macrophage labeling revealed the regular structure of the hepatocytes and the clear appearance of three liver macrophages in this field of view (Figure 2a). After segmentation and reconstruction in Imaris, the hepatocyte cell walls were represented both as 3D surfaces and as Cells objects (Figure 2b), from which the membrane were extracted and represented as a confluent cell layer (Figure 2c). The latter representation allowed the enclosed cell surfaces to be tested for bacterial content. Here the number of bacteria was determined by counting the engulfed Spots (uniform-diameter green spheres from Imaris in Figure 2c) per macrophage, used later to characterize the polarization-dependence of *S. aureus* uptake by M0, M1 and M2 macrophages. The distribution of high bacterial—content macrophages was not uniform, as shown by the clustering of yellow-to-red color-coded cells in Figure 2c.

The endothelial and hepatic sides of the chips were identified based on the Z position of the segmented cells relative to the axial location of the scaffolding membrane, as shown in Figure 1b. The membrane itself was not labelled, but it was possible to precisely identify its location by applying contrast stretching to the green *S. aureus* channel, inverting it, and then using the resulting bright disks that appeared in a narrow Z range in the Z slice signal to generate Spot objects that corresponded to the membrane pores (Figure 3a-b). This approach allowed a simple way to get a 3D map of all the membrane pores, thus precisely determining the side at which a given cell is located (Figure 3c-d). In the example in Figure 3c-d, three macrophages were identified and segmented using the lateral view (Figure 3c). When compared with the localization of the previously identified membrane in axial view, it became clear that two of three macrophages were located on the hepatic side of the chip, whereas the third immune cell was on the vascular side; all three macrophages were located close to the membrane (Figure 3d).

As the *S. aureus* cells were visualized as Spots after segmentation, the spatial location of the intensity center of the bacteria was determined precisely, whereas information about the exact shape of the pathogens was lost (Figure 3e; for technical details on how the Imaris parameters were set for the Spots and Surfaces objects and how these were utilized to measure infection, see [24]. This was a meaningful compromise, because the necessary optical resolution to observe the shape of individual bacteria was not available with the technology applied in this work. The segmented Spots were categorized according to their position: inside the endo- or epithelial cells or the macrophages, adherent to the cell membranes, or dissociated (Supplementary Figure 2 bottom right, Figure 3e). The macrophage and the *S. aureus* in the example in Figure 3e illustrate how a typical spatial distribution of these cells look like: the majority of the bacteria were already phagocytosed by the macrophages (these are the magenta dots in Figure 3e), with a minority being attached to the macrophage surface (adherent pathogens, see the cyan spots in Figure 3e). The rest of the bacteria were classified as dissociated, i.e., being more than one micrometer away from any macrophage surface (see the silver spots in Figure 3e). This elaborate classification technique was utilized to quantify the number of intracellular bacteria when faced with immune cells of various polarization states.

The assembly of these novel image quantification workflows was utilized to extract information from the 3D microscopy images taken 2h p.i. about the efficiency of macrophages in clearing out bacterial pathogens, the roles of endothelial and parenchymal cells in taking up bacteria, and the role of macrophage polarization in *S. aureus* uptake. Firstly, we could demonstrate the major role of macrophages in clearing *S. aureus* from the circulation in the liver model. Similar to *in vivo*, macrophages had a protective role by preventing the infection of endothelial cells and hepatocytes with *S. aureus*, which did not depend on the macrophage polarization pattern (Figure 4a, b). However, in the absence of macrophages, considerable high numbers of intracellular bacteria were detected within endothelial cells as well as hepatocytes. The vast majority of bacteria was detected in macrophages, with the highest bacterial counts found in M2 polarized macrophages 2 h p.i. as well as 48 h p.i. (Figure 4 c,d). The MOI was kept constant at the time of infection regardless of macrophage polarization, still bacterial counts were higher in M2 macrophages throughout the course of infection. Thus, the initial bacterial uptake rather the ability to differentially clear persisting bacteria among macrophage polarization conditions might determine the intracellular bacterial load 48h p.i.. Although not statistically significant, a similar trend was observed in 2D macrophage monocultures under static conditions (Supplementary Figure 1b). We next studied the capability of *S. aureus* to disseminate after intracellular persistence by analyzing planktonic bacteria at the vascular and parenchymal side effluent (Supplementary Figure d). Dissemination of *S. aureus* was primarily seen at the parenchymal side of the model. The polarization pattern of macrophages had no significant impact on the dissemination rate, with only a few bacteria escaping from the liver model (Figure 4e, f). The analysis of the formation of SCVs revealed a positive correlation with the total amount of persisting bacteria 48 h p.i.. Importantly, SCV formation was not observed in the 2D mono-cell culture of macrophages, indicating the contribution of other cell types in this process (Figure 4g).

**Figure 4.**
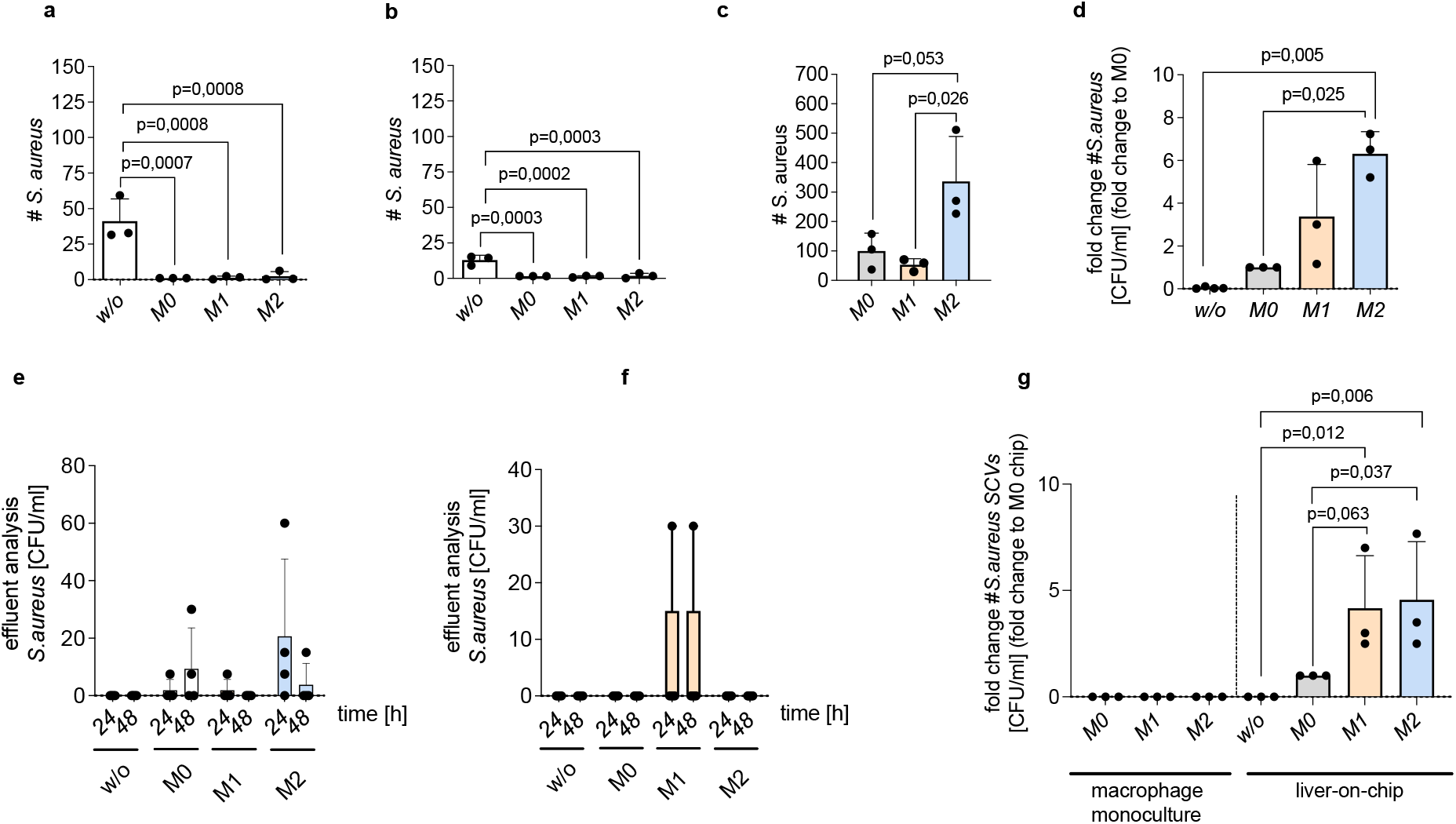
Uptake, persistence, dissemination, and SCV formation of S. aureus in liver-on-chip. a-c) image analysis-based quantification of the number of persisting S. aureus per region of interest (ROI) in a) endothelial cells; in b) hepatocytes or in c) macrophages of liver-on-chip without cocultured macrophages (w/o) or in coculture with non-activated (M0), M1 or M2 polarized macrophages 2 h p.i.. At least 5 randomly selected ROIs per condition were analyzed. d-g) CFU counts 48 h p.i. of d) lysed liver-on-chip; e-f) disseminated S. aureus within e) the parenchymal effluent, f) the vascular effluent. g) CFU analysis of SCVs formed in macrophage monocultures or liver-on-chip (SCVs identified based on colony size (Supp. Fig. 1c)). d, g) CFU counts were normalized as fold changes to the values of the M0 condition per experiment and macrophage donor. a-g) Bars indicate the mean of values from three independent experiments, error bars indicate SD. Statistical analysis shows p-values of one-way ANOVA testing.

Cytokine release profiles within the effluent from liver-on-chip validated an acute immune response to *S. aureus* infection with IL-1β levels increasing at 24 and 48 h p.i. in liver-on-chip containing macrophages. Similar trends were observed for the release of the cytokines IL-18, IL-6, TNFα, and IL-10 (Figure 5a-e). Except for the release of IL-6, where baseline cytokine levels were already detectable without cocultured macrophages, the immune response to *S. aureus* infection strongly depended on the presence of macrophages. Still, the initial macrophage polarization pattern at the time of infection had no significant effect on cytokine release. These results suggest that pre-infection macrophage polarization determines the number of *S. aureus* capable of infecting the cell but does not influence the immune response to intracellular persisting bacteria. Further, in liver-on-chip with M2 macrophages we observed a significant increase of lactate dehydrogenase (LDH) and aspartate aminotransferase (ASAT) release into the effluent suggesting loss of cellular integrity that was associated with a drop of the albumin synthesis rate 48h p.i., reflecting reduced hepatocyte metabolism (Figure 6a-c).

**Figure 5.**
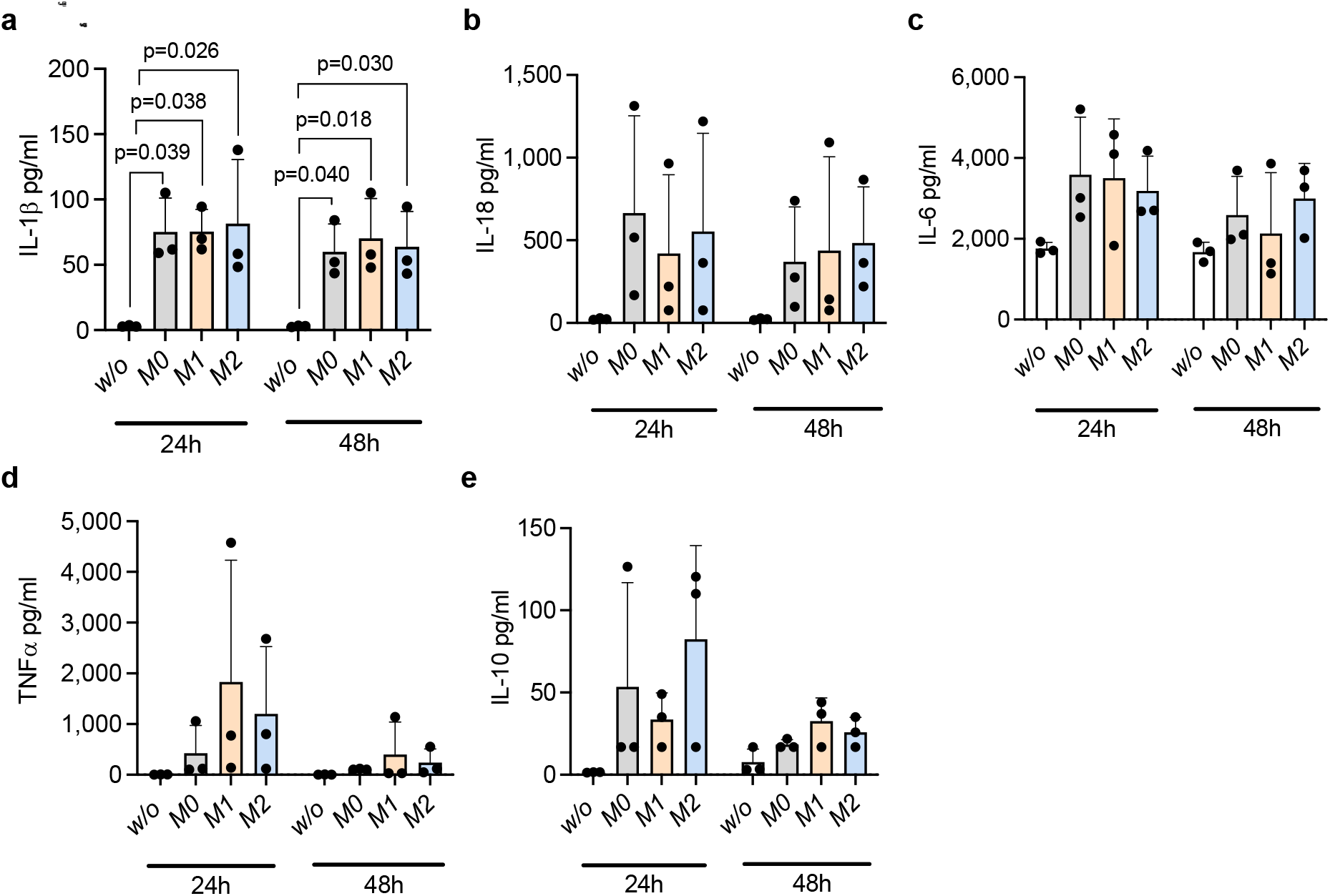
Cytokine profiles measured 24 or 48 h p.i in the vascular effluent of liver-on-chip without cocultured macrophages (w/o) or in the presence of non-activated (M0), M1 polarized, or M2 polarized macrophages. a-e) Bars indicate the mean of values from three independent experiments, error bars indicate SD. Statistical analysis shows p-values of one-way ANOVA testing.

**Figure 6.**
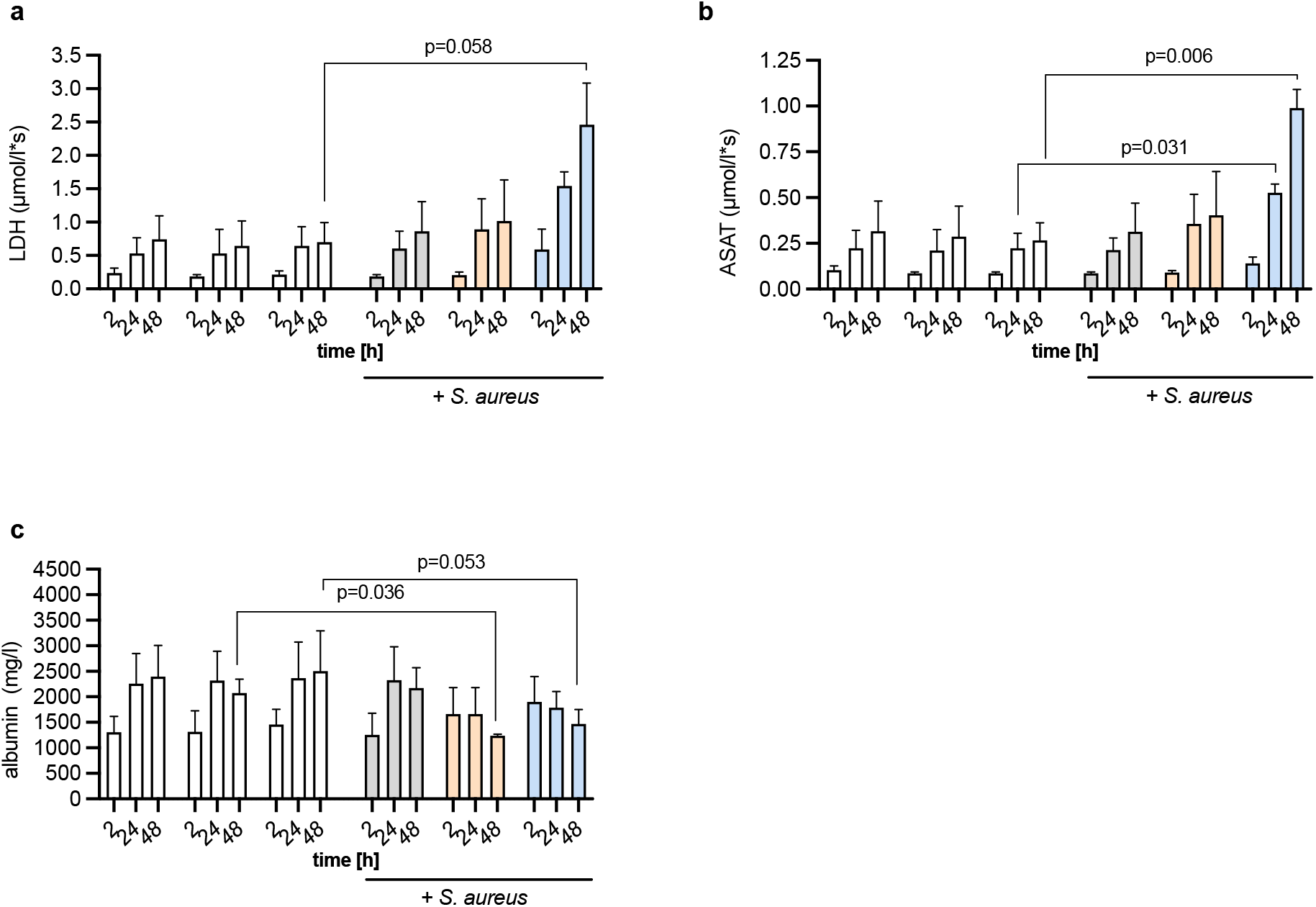
Effluent analysis of the release of lactate dehydrogenase (LDH), aspartate aminotransferase (ASAT), or albumin at the parenchymal side of the liver-on-chip model 48 h p.i.. a-c) Bars indicate the mean values from three independent experiments, error bars indicate SD. Statistical analysis shows p-values of one-way ANOVA testing.

We next analyzed macrophage counts in liver-on-chip after infection. No differences were observed 2 h p.i. with respect to macrophage polarization stages. However, a declining number of M2 polarized macrophages was detected 48 h p.i. in infected liver-on-chip (Figure 7a, b). We then asked whether the macrophage loss as a consequence of infection could be compensated by recruiting monocytes from the circulation. Monocyte perfusion experiments proved an increased monocyte adhesion rate upon infection compared to non-infected liver-on-chip (Figure 7c). This observation underlines the importance of macrophages in orchestrating monocyte infiltration during infection with *S. aureus*. In particular, in the presence of M0 macrophages, monocyte counts infiltrated in the liver model were elevated 48 h p.i. compared to liver-on-chip without macrophages. Interestingly, monocyte recruitment was diminished to basal levels in the presence of infected M2 macrophages similar to liver-on-chip without cocultured macrophages (Figure 7c, d). This finding indicates that in the presence of M2 macrophages, the ability to replenish macrophage loss upon infection is disturbed. We found no indication of bacterial uptake by recruited monocytes at sites of infection, where intracellular presence of *S. aureus* was strictly restricted to macrophages (Figure 7e). Further, we did not detect viable bacteria by CFU assays in monocytes from the effluent of infected liver-on-chip (data not shown).

**Figure 7.**
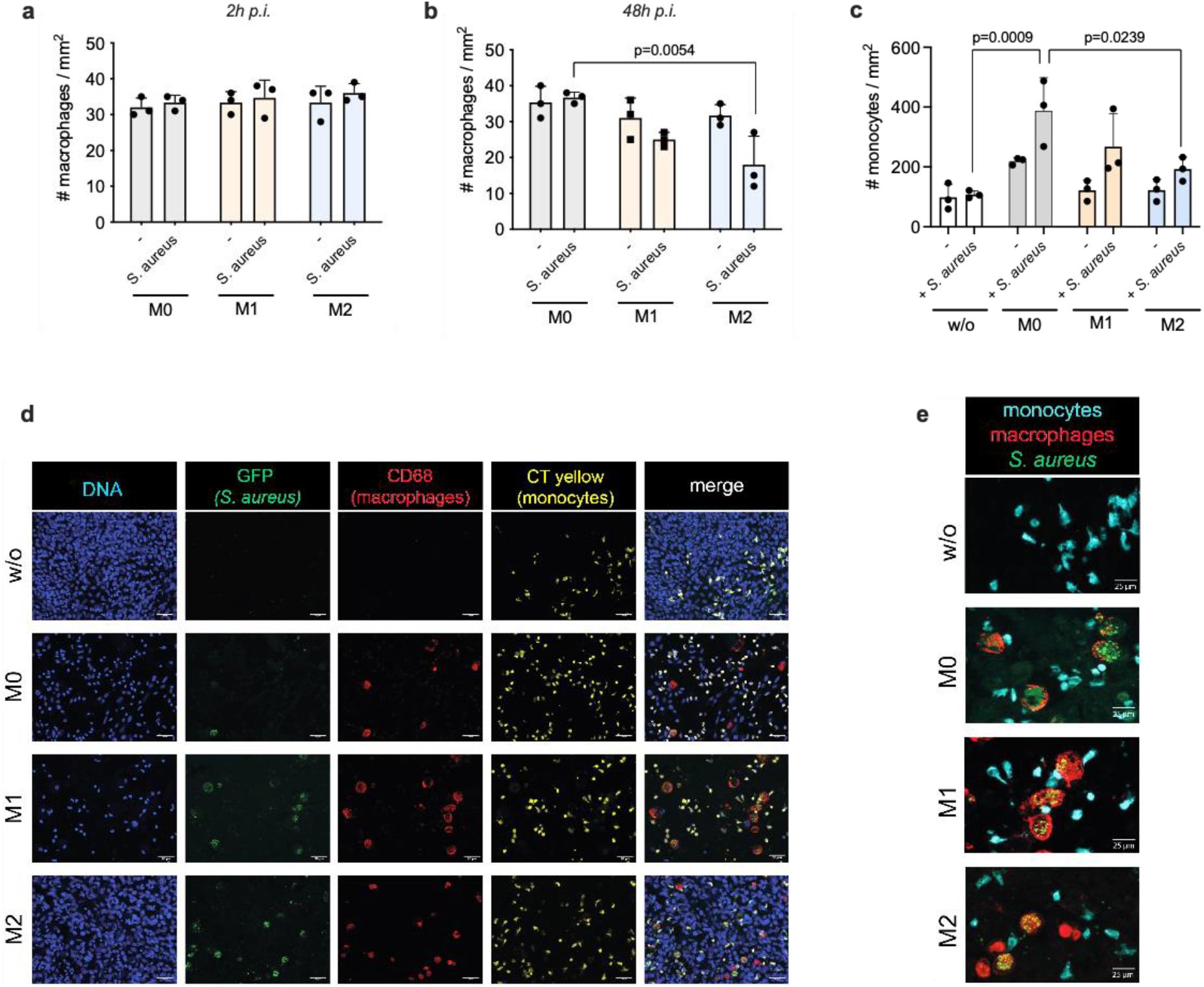
Macrophages and monocytes count in liver-on-chip. a-b) Number of macrophages per mm^2^ in liver-on-chip a) 2 h p.i. or b) 48 h p.i.; c) number of recruited monocytes per mm^2^ 48 h p.i.. a-c) Image analysis-based quantification of five randomly selected RIO per condition. d-e) Representative images of monocyte recruitment experiments d) scale bar 50 μm; e) scale bar 20 μm. a-c) Bars indicate mean values from three independent experiments, error bars indicate SD. Statistical analysis shows p-values of one-way ANOVA testing.

## Discussion

Macrophages play a central role in removing planktonic *S. aureus* from the bloodstream by the liver and kill bacteria after uptake by phagocytosis. For the first time, we were able to leverage a complex human liver-on-chip model with tailored image analysis procedures to follow these processes in vitro and demonstrate the prominent role of macrophages and its polarization on facilitating intracellular persistence and metabolic adaption of *S. aureus* in a protected immunological niche.

Our results show that increased numbers of bacteria initially phagocytosed by macrophages result in higher amounts of living persisting bacteria capable of surviving by exploiting this cell type as a cellular niche, likely by exhaustion of antimicrobial functionality over time. This mechanisms contribute to the strategy of the pathogen to remain undetected by peripheral immune cells such as neutrophils [6d]. The limited capacity of intracellular killing by macrophages is the rate-limiting process in bacterial clearance. In this context, disturbed maturation of the phagosome and a decreased acidification with reduced activation of cathepsin D have been discussed to be associated with the failure of inducing apoptosis-associated bacterial killing in macrophages [11b]. Consequently, viable bacteria load further increases by ongoing phagocytosis and form a pool of adapted variants capable of lysing and reinfecting other macrophages [11b]. A recent study showed that also replacing the majority of the infectious burden with commensals could augment infection and trigger the formation of “proinfectious agents” as a result of finite macrophage clearing capacity [25]. In mice, this mechanism has been proven to contribute to the dissemination into other organs in the periphery, such as the kidneys [6d].

Recent studies revealed that various gram-negative and gram-positive bacteria induce similar transcriptional activity termed “core response to infection,” related to M1 polarization [26]. However, other studies indicate that macrophages could respond to *S. aureus* exposure differently, promoting pro-inflammatory M1 as well as anti-inflammatory M2 macrophage responses [27]. Experimentally, M1 or M2 polarization is often induced *in vitro* by cytokine stimulation of macrophage monocultures. However, single-cell type models neglect the complex microenvironment and the contribution of other cell types in the course of infection. Thus, macrophage monocultures might not entirely correlate to phenotypes found *in vivo* that constantly change over time depending on the tissue context and the stage of infection [28]. Macrophage plasticity is very rapid and enables the cell to shape a dynamic immunomodulatory environment. The cells orchestrate the initial phase of inflammation and set the stage for later phases of infection with the resolution of inflammation and subsequent tissue remodeling [29]. Thus, i*n vivo* macrophage activation is delineated as a continuum composed of multiple transient functional states between M1 and M2 in which the cell reversely adapts to its environment, which is formed by released growth factors and chemokines during acute infection and tissue repair [30]. In order to limit detrimental effects to the host tissue, an M1 to M2 polarization shift of macrophages is induced during sepsis in the transition from the acute “systemic inflammatory response syndrome” (SIRS) to the “compensatory anti-inflammatory response syndrome” (CARS) associated with immune tolerance to pathogens [31]. M2 macrophages have a potent capacity of phagocytosing cell debris, apoptotic and dead cells. Still, they are poor antigen-presenting cells which relates to their function in limiting the pro-inflammatory host response [32]. Patients suffering from chronic rhinosinusitis with nasal polyps show an increased *S. aureus* colonization of the nasal mucosa associated with an M2 activation phenotype that is linked to decreased phagocytosis and an increased bacterial persistence [33]. Several studies demonstrated the ability of different pathogens to manipulate macrophage activation to neutralize antimicrobial effectors and to establish chronic infections. Intracellular persisting bacterial pathogens including *Mycobacterium tuberculosis, Salmonella, Coxiella burnetii*, and others have been reported to manipulate macrophage polarization, including reprogramming of macrophage polarization from M1 to M2 as a strategy to persist and to evade immune recognition, which favors the onset of chronic infections [26a, 32, 34]. In our study, we could prove that macrophage polarization at the time of infection determines the load of intracellular bacteria already 2 h p.i.. Recent studies show that macrophages take up *S. aureus* via classical endocytosis within 1.5 h [6d]. In our human liver model, we found significantly higher numbers of viable intracellular bacteria in M2 polarized macrophages 48 h p.i.. This is likely an effect of the increased phagocytosis rate of M2 macrophages, which represents a hallmark of this macrophage phenotype [35]. In the liver model, increased numbers of M2 macrophage-persisting bacteria were associated with a significant loss of viable macrophages 48 h p.i. and increased cell death with reduced metabolic activity of hepatocytes, probably as a consequence of hepatocyte impairment by damage-associated molecular patterns released from dying macrophages.

*In vivo*, the loss of Kupffer cells led to a higher susceptibility and severity of *S. aureus* infections [12a]. Several reports have demonstrated that following *S. aureus* infection, monocytes are recruited to sites of infection as a result of an up-regulation of various cell adhesion molecules expressed by endothelial cells [36]. Increased monocyte recruitment is a mechanism to replenish macrophage loss and is important in the course of infection for efficient eradication of the pathogen. Significant recruitment of monocytes was observed in liver-on-chip models containing non-activated macrophages but was significantly impaired in the presence of M2 polarized macrophages with higher bacterial loads. This highlights the role of functional macrophages in orchestrating monocyte recruitment in the course of infection and might further point to a mechanism that would allow *S. aureus* to prevent its eradication from the tissue by restricting monocyte infiltration and macrophage differentiation at sites of infection. Recently, it has been demonstrated in mice receiving intracellular viable *S. aureus* by injection of infected bone marrow-derived macrophages that these animals had a higher bacterial load in the kidney and liver compared to mice that received injection of similar amounts of planktonic *S. aureus* [37]. However, we found no viable bacteria been ingested by monocytes that migrated into the cell layers or remained within the effluent.

Although bacterial burden was significantly elevated in liver-on-chip with M2 macrophages, we found no indication of elevated dissemination into the effluent. Recently, animal models have shown that a small population of bacteria is enriched by recurring infection and cell lysis cycles of macrophages. This “population bottleneck” proposed by Pollitt et al. argues for the accumulation of selected variants fitter and better adapted to the specific microenvironment rather than the formation of new bacterial mutants [12a]. Similar effects might be responsible for the accumulation of bacteria within our liver model, where the dissemination of bacteria strictly relied on the presence of functional macrophages. We did not observe a correlation of increasing bacterial counts within macrophages with elevated release into the effluent. This fact might be considered a hint that persisting *S. aureus*, after host cell lysis, prefers to jump to other macrophages within the tissue due to a variant enrichment rather than disseminating into the effluent. Targeting such an adapted variant population of bacteria by tailored antimicrobial treatments will likely help to restrict bacterial dissemination and limit the relapsing capacity of *S. aureus* in chronic infections [11b]. Follow-up studies will look at more extended circulation and migration times of monocytes which would allow its differentiation to macrophages and will also address the role of neutrophil granulocytes as potential “trojan horses” for *S. aureus* to systemically disseminate to other organs [38].

A common problem in treating macrophage persisting *S. aureus* variants is the fact that the antibiotic of choice, vancomycin, which is frequently used to eradicate MRSA infections, cannot penetrate the membrane of macrophages to kill persisting bacteria [9c]. The problem in treating these infections is further complicated by the ability of *S. aureus* to enter the dormant SCV phenotype characterized by a reduced metabolic activity which significantly limits the ability of the antibiotic to interfere with bacterial survival [13, 39]. SCV formation has been reported from the USA300 and other strains as a strategy to survive in abscesses by upregulation of adhesin expression and downregulation of bacterial metabolism with reduced expression of virulence factors facilitating an improved intracellular persistence and the evasion of immune cell detection [13, 39]. Further, SCVs could prevent induction of hypoxia-inducible factor, an important sensor for intracellular persisting pathogens, and thereby circumvent an appropriate antimicrobial response favoring abscess formation [40]. We observed a higher number of SCVs due to increased bacterial loads within macrophages cocultured in liver-on-chip. Interestingly, SCV formation was not observed in macrophage monocultures obtained from the same donors. This agrees with reports from another group reporting no elevated SCV formation in macrophages differentiated from THP-1 cells by phorbol 12-myristate 13-acetate stimulation within five days in a mono-cell culture [41]. Further, monocyte-derived macrophages in mono-cultures do not undergo induction of apoptosis or necrosis during *S. aureus* persistence and remain functional for several days after stimulation [6d]. Although we obtained similar results 48 p.i. of macrophages in the monoculture, we overserved significant cell loss of M2 macrophages in liver-on-chip associated with increased formation of SCVs. These results imply that signals from additional cell types are required to create a milieu supportive for S. aureus persistence within M2 macrophages.

The translational relevance of our findings is underlined by several studies reporting that *S. aureus* infections are an important clinical complication in patients with chronic liver diseases such as fibrosis. In these patients, bacteremia is the most common type of *S. aureus* infection and is associated with a high mortality rate that is higher compared to any other disease coinciding with a *S. aureus* infection [42]. In particular, cirrhosis has been identified as an independent risk factor for bloodstream infections in patients with hospital-acquired pneumonia caused by *S. aureus* [43]. Chronic liver disease is often linked to higher counts of M2 polarized macrophages in the liver [44] that, according to our study, have impaired bacterial clearance capacity, which results in higher bacterial loads and increased frequency of SCV phenotype switching. This adaption by the pathogen to the microenvironment under infection conditions enables it to hide from immune cell detection and limits the host’s options to eradicate persisting infections. In addition, the impaired replenishment of macrophages by circulating monocytes observed *in vitro* after the infection-related loss of M2 macrophages may contribute to bacterial persistence within the liver *in vivo*. Our study’s limitation consists of the limited number of donors due to challenging logistics in cell sourcing of primary monocytes and the need to use different cell donors for tissue engineering of the respective cell types to assemble liver-on-chip models. Future studies should leverage human-induced pluripotent cells as a cellular source of individual cell types to create isogenic tissue models that reflect individual donor-specific genetic backgrounds [45]. In this proof-of-concept study, we could demonstrate for the first time the potential of liver-on-chip to study bacterial infection and persistence mechanisms in macrophages. Future studies with advanced microphysiological models will allow higher sample numbers to provide a more detailed insight into the mechanisms of adaption of *S. aureus* to environmental cues and its strategy to persist and disseminate within the human liver.

## Author Contributions

F.S., S.C., M.G., A.S. performed experiments and analyzed data. Z.C. and M.T.F. developed strategies and implemented algorithms for the image-based quantification of *S. aureus*. B.G.J.S. provided GFP-expressing *S. aureus*. M.T.F., B.L., L.T., O.W., and A.S.M. planned and supervised experiments. A.S.M. designed the study and wrote the manuscript. F.S., Z.C., S.C., M.G., A.S., B.G.J.S., O.W., M.T.F., B.L., L.T., A.S.M critically revised the manuscript.

## Author Contributions

The Deutsche Forschungsgemeinschaft financially supported this work under grant number MO 2968/1-1 to A.S.M. and grant number 1618/5-1 to B.L., and by the Cluster of Excellence “Balance of the Microverse” under Germany’s Excellence Strategy - EXC 2051 - Project-ID 690 390713860 to M.T.F., O.W., B.L., L.T., and A.S.M. The work was further financially supported by the Collaborative Research Center PolyTarget 1278 (project number 316213987) to M.T.F., O.W. and A.S.M. and the German Ministry for Education and Research (BMBF) under grant no. 01EO1002 (CSCC) to B.L., L.T., and A.S.M; and grant no. 01EC1901B (MESINFLAME) to B.L. and L.T.

