## Supplementary material for "Human macrophage polarization determines bacterial persistence of *Staphylococcus aureus* in a liver-on-chip-based infection model": Supplemetal Figures 1 & 2

### Supplementary Information

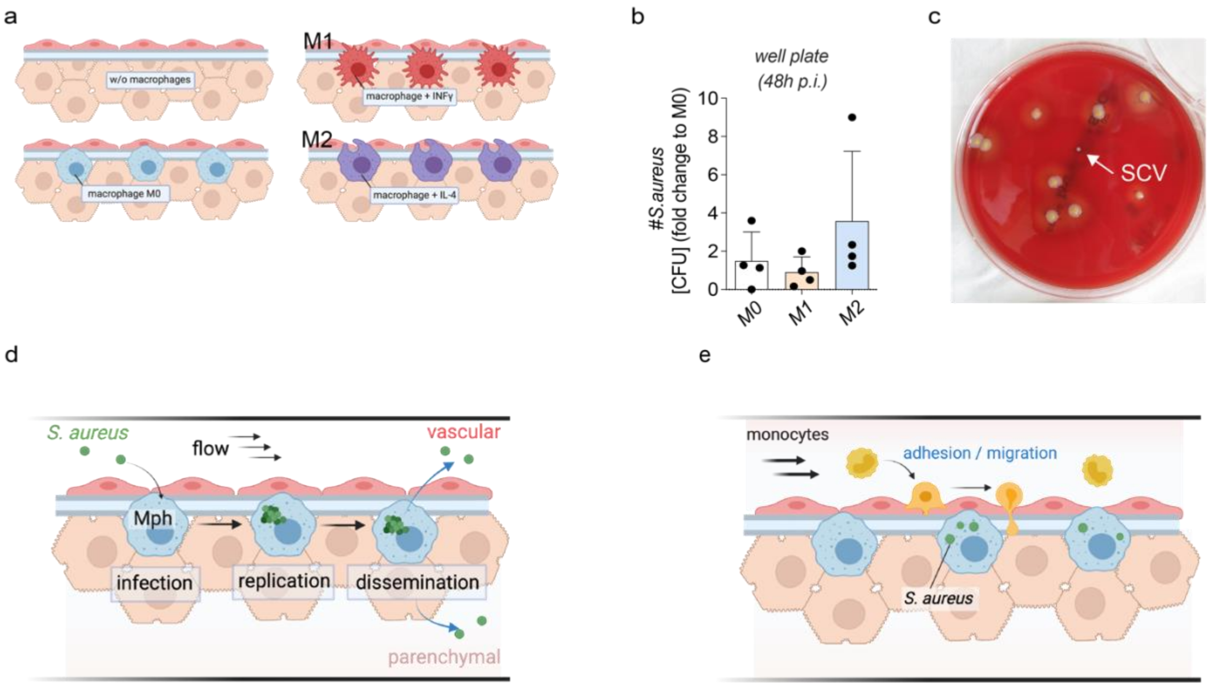


**Supplementary Figure 1**. *a) Testing conditions in liver-on-chip for the role of macrophages and its activation/polarization stages as a niche in persistence, phenotype switching, and dissemination of S. aureus; b) number of CFU from S. aureus infected monocytes 48h p.i. in macrophage monocultures. Bars indicate the mean; error bars indicate SD. Data of three independent experiments is shown. c) representative image of SCV colony (white arrow points to SCV) cultured on a sheep blood agar plate. d) scheme of S. aureus infection of macrophages (Mph), intracellular replication and dissemination after macrophage lysis in perfused liver-on-chip. e) Scheme of monocyte recruitment assays in perfused liver-on-chip.*

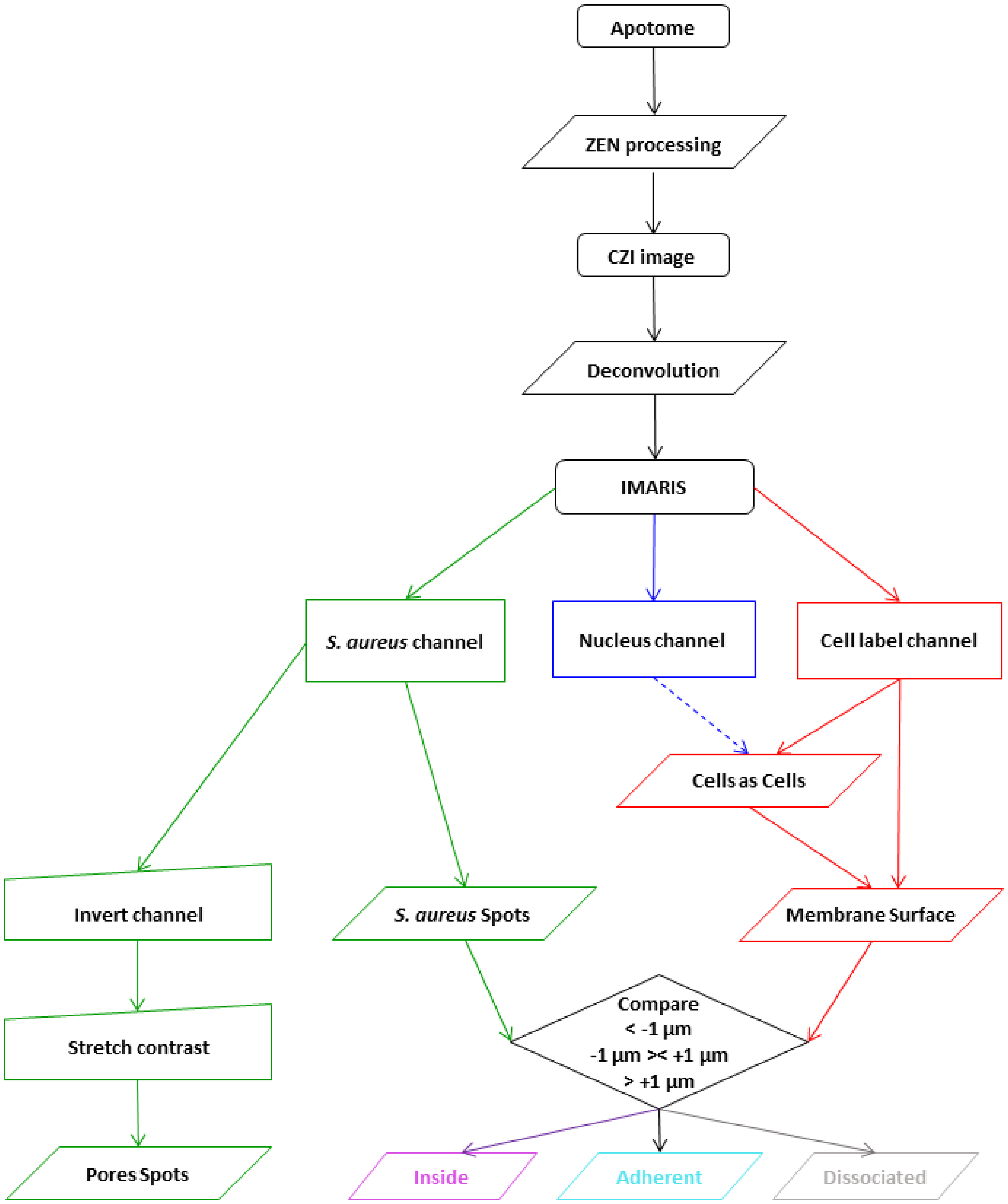


**Supplementary Figure 2**. *The workflow of image quantification applied in the organ-on-chip analysis. The original images were preprocessed in Zeiss ZEN in order to calculate the deblurred images from the raw Apotome images series, followed by deconvolution of the CZI images in Huygens Professional (lines and flowchart elements in black) and then processing in Imaris. Here the three fluorescence channels were separated (the transmitted light channel was generally ignored during the rest of the analysis) into bacterial (green), endo/epithelial/macrophage (red) and nuclear (blue) labelling. The nuclear and cell labelling signals were used to segment the cells as Cells objects; alternatively, the membrane signal alone was used for the Cells generation (the blue dashed line indicates that the nuclear signal*

*was not always utilized). The membrane surface was produced by exporting the membrane component of the Cells object in Imaris. The S. aureus cells were identified from the corresponding green fluorescence channel (lines and flowchart elements in green). The green channel was also used to identify the membrane pores by inverting the bacterial channel, stretching its contrast, and segmenting the pores as Spots based on the bright disks that appeared in the narrow Z zone where the membrane was located (bottom left segment of the flowchart; see also Supplementary Figures 4a-b). The Spot objects identifying S. aureus cells were categorized according to their distance from the corresponding cell membrane (macrophage; endothelial cell; hepatocyte): the Spots below -1 µm from the cell membrane were classified as being inside the cells (bottom right, magenta); those between -1 µm and +1 µm were identified as adherent bacteria (bottom right, cyan), whereas those farther than +1 µm from the membrane were considered dissociated (bottom right, silver).*
